# *FKS1* is required for *Cryptococcus neoformans* fitness *in vivo*: application of copper-regulated gene expression to mouse models of cryptococcosis

**DOI:** 10.1101/2022.03.24.485727

**Authors:** Sarah R. Beattie, Andrew J. Jezewski, Laura C. Ristow, Melanie Wellington, Damian J. Krysan

## Abstract

There is an urgent need for new antifungals to treat cryptococcal meningoencephalitis, a leading cause of mortality in people living with HIV/AIDS. An important aspect of antifungal drug development is the validation of targets to determine whether they are required for the survival of the organism in animal models of disease. In *Cryptococcus neoformans*, a copper-regulated promoter (pCTR4-2) has been used to modulate gene expression *in vivo* previously. The premise for these experiments is that copper concentrations vary depending on the host niche. Here, we directly test this premise and confirm that the expression of *CTR4*, the promoter used to regulate gene expression, is much lower in the mouse lung compared to the brain. To further explore this approach, we applied to the gene encoding 1,3-β-glucan synthase, *FKS1*. *In vitro*, reduced expression of *FKS1* has little effect on growth but does activate the cell wall integrity stress response and increase susceptibility to caspofungin, a direct inhibitor of Fks1. These data suggest that compensatory pathways that reduce *C. neoformans* resistance do so through post-transcriptional effects. *In vivo*, however, a less pronounced reduction in *FKS1* expression leads to a much more significant reduction in lung fungal burden (~1 log_10_ CFU), indicating that the compensatory responses to a reduction in *FKS1* expression are not as effective *in vivo* as they are *in vitro*. In summary, use of copper-regulated expression of putative drug targets *in vitro* and *in vivo* can provide insights into the biological consequences of reduced activity of the target during infection.

**Importance:** Conditional expression systems are widely used to genetically validate antifungal drug targets in mouse models of infection. Copper-regulated expression using the promoter of the *CTR4* gene has been sporadically used for this purpose in *C. neoformans*. Here, we show that *CTR4* expression is low in the lung and high in the brain, establishing the basic premise behind this approach. We applied it to the study of *FKS1*, the gene encoding for the target of the echinocandin class of 1,3-β-glucan synthase inhibitors. Our *in vitro* and *in vivo* studies indicate that *C. neoformans* tolerates extremely low levels of *FKS1* expression. This observation provides a potential explanation for the poor activity of 1,3-β-glucan synthase inhibitors toward *C. neoformans*.

## Introduction

*Cryptococcus* spp. are one of the most important human fungal pathogens with global effects on human health, particularly for people living with HIV/AIDS (1, 2). Serological surveillance suggest that most people have been exposed to *Cryptococcus* early in life (3). For the vast majority, this exposure does not lead to disease. However, individuals with altered T-cell function and, indeed, some with apparently normal immune dysfunction develop cryptococcal meningoencephalitis (CME). CME is uniformly fatal unless treated and is a leading cause of death for people living with HIV/AIDS (4). Importantly, CME can be the sentinel event leading to the diagnosis of HIV infection. As a result, many people must first survive CME to take advantage of the lifesaving advances in the treatment of HIV.

The first line regimens for the treatment of cryptococcal meningitis are currently based on various combinations of amphotericin B, flucytosine, and/or fluconazole (5). Unfortunately, these medications all have issues that limit their effectiveness and/or utility. Amphotericin B is toxic, requires intravenous administration, and necessitates laboratory monitoring of electrolytes (6); these characteristics can make amphotericin logistically difficult to use in resource-limited regions without extensive medical infrastructure. Flucytosine is also toxic and is not available in many countries that have the highest rates of disease (7). Finally, fluconazole is cheap and available but its efficacy is poor as a single agent and must combined with either amphotericin B or flucytosine in order to have a reasonable outcome. The development of safe and effective therapies for CME that are accessible to people who live in the areas with the highest rates of disease should be a high priority for modern medicine.

To address the unmet clinical need for new anti-cryptococcal therapies, a number of groups have explored repurposing and new chemical entity discovery approaches based on both high throughput phenotypic screening and target-focused medicinal chemistry campaigns (8). A key task during pre-clinical drug development is to validate that a putative target leads to reduced growth or virulence of the fungus in an animal model of infection. Usually, these targets are encoded by genes that are essential for viability making genetic studies more complicated, particularly in assessing the contribution to virulence. The most common approach to evaluate essential genes in models of fungal infection is the construction of conditional expression alleles that are responsive to tetracycline/doxycycline: the so-called Tet_OFF_ technology (9–11). A major advantage to this system is the ability to administer doxycycline to mice which can then regulate fungal gene expression *in vivo* (12–14). To date, this technology has not been applicable to *C. neoformans* (15). Modulation of some genes has been achieved but it requires extremely high doses of doxycycline that are not achievable in mice. The reasons for this technical limitation are not clear but similar problems have been reported for the application of doxycycline regulation to *Ustilago maydis*, another basidiomycete (16). Thus, in *C. neoformans*, genetic validation studies have been largely limited to either *in vitro* studies using alternative conditional promoters or *in vivo* analysis of non-essential genes.

One of the most widely used system for the conditional expression of genes in *C. neoformans* is based on the promoter for the copper transporter, *CTR4*. Initially developed by the Doering lab, replacement of the promoter for the gene of interest with multiple copies of the *CTR4* (p*CTR4-2*) leads to an allele that is highly expressed in low copper and repressed in the presence of high concentrations of copper (17). Prior to our recent work (18), there had been one report of using a *CTR4*-regulated gene to assess gene function in the setting of a mouse model of cryptococcal infection. In this work, the promoters of the essential fatty acid synthases *FAS1*/*FAS2* were replaced by p*CTR4-2* (15). *In vitro*, neither strain was able to grow in the presence of 25 μM copper sulfate while only the p*CTR4-2-FAS2* strain showed reduced fungal burden in the lungs compared to the H99 following intranasal infection. No dissemination to the brain was detected with the p*CTR4-2-FAS2* strain while the p*CTR4-2-FAS1* showed a ~1 log_10_ reduction in brain burden. These observations are consistent with the likelihood that cryptococcus cells in the lung experience a relatively high copper concentration, leading to reduced expression of *FAS2* and reduced fungal burden. In addition, the discordance between the *in vitro* and *in vivo* phenotypes of the p*CTR4*-2-*FAS1* strain suggest that the effect of *pCTR4-2* regulation of gene expression may vary with the specific gene under regulation.

Recently, we reported that infection of mice with a strain that contained *HSP90* under control of p*CTR4-2* lead to reduced expression of *HSP90* in the lung relative to the congenic control whereas the same strain inoculated intravenously and sampled from the brain had dramatically increased expression relative to control (18). Consistent with this expression data, mice infected intranasally with the p*CTR4-2-HSP90* strain survived longer than those infected with the wild type and there was no difference in survival between the two strains in the intravenous inoculation experiment. These data indicate that in *C. neoformans* experiences high copper conditions in the lung and low copper conditions in the brain and that p*CTR4-2*-regulated alleles might be useful to validate drug targets or study essential gene function in mouse models of cryptococcosis. Here, we directly measure the expression of *C. neoformans CTR4* in infected lung and brain and confirm that it is much lower in the lung as compared to the brain, supporting the previous observations and the utility of this approach for some genes.

To further explore the use of p*CTR4-2* in the regulation of essential genes related to antifungal therapy, we generated a *pCTR4-2-FKS1* strain and examined it’s in vitro and in vivo phenotypes. Fks1 is a 1,3-β-glucan synthase and the target of the echinocandin class of antifungal drugs (19). Echinocandins such as caspofungin have poor activity against *C. neoformans* and are ineffective in mouse models of infection (20). This in effectiveness is despite genetic evidence indicating that *FKS1* is an essential gene (21) and biochemical experiments indicating that Fks1 is inhibited by echinocandins (22). The mechanism of echinocandin ineffectiveness against *C. neoformans* is unclear and remains an area of active investigation. Here we show that *in vitro*, *C. neoformans* tolerates drastic reductions in *FKS1* expression with minimal changes in growth; a modest increase in echinocandin susceptibility and activation of the cell wall integrity response. *In vivo*, the fitness of a *pCTR4-2-FKS1* strain is reduced in the lung to a greater extent than observed in vitro. These data suggest that *C. neoformans* is remarkably resistant to reductions Fks1 activity *in vitro* but somewhat more susceptible *in vivo*.

## Results

### *C. neoformans* cells infecting the brain have elevated *CTR4* expression relative to cells infecting the lung

Pioneering studies of copper homeostasis in *C. neoformans* during infection by Waterman *et al*. indicated that *C. neoformans* copper is limiting during infection of macrophages and the CNS but not during pulmonary infection (23). Ding *et al*. used bioluminescence to show that expression of the metallothionein *CMT1*, which is induced during high copper, is elevated in the lung relative while *CTR4* is detectable but relatively lowly expressed (24). The elevated ratio of *CMT1*/*CTR4* is consistent with *C. neoformans* occupying a niche in the lung that is relatively copper replete. Further supporting that conclusion, Waterman *et al*. also found that deletion of *CUF1*, the gene encoding a transcription factor that activates *CTR4* expression, had no effect on replication of *C. neoformans* in the lung but prevented dissemination to the brain (23). Although *CTR4* expression appears to differ between *C. neoformans* infecting the lung and the brain of mice, it is required for virulence in both pulmonary and intravenous inoculation models. The differential expression data suggested that it might be possible to use *CTR4-2*-regulated alleles as knockdown alleles to study essential or severely compromised mutants during pulmonary infection; alternatively, during CNS infection the construct could display features of overexpression.

To our knowledge, the expression of *CTR4* in *C. neoformans* during infection of mouse lung or brain has not been directly characterized using RT-PCR; previous data were derived from experiments using either fluorescent or luminescent *CTR4*-fusion reporters. To directly measure *CTR4* message *in vivo*, we inoculated AJ/cr mice with the reference strain KN99α using the intranasal and the intravenous inoculation route; to reiterate, the intravenous route establishes CNS and pulmonary infection within hours while the intranasal route establishes pulmonary infection immediately but can take over a week to disseminate to the brain. We harvested lungs and brains from animals inoculated by each route on days 4- and 8-post inoculation. The expression of *CTR4* in the lungs was the same between the intravenous and intranasal models; on day 4 expression was low and increased slightly by day 8 in both models (Figure 1). We did not detect any *CTR4* or *TEF1* expression in either 4- or 8-dpi brain samples from the intranasal model, suggesting that these timepoints occur before significant dissemination to the brain. In the brains collected from intravenously-inoculated animals, we observed much higher expression of *CTR4*. Compared to the lungs, the expression of *CTR4* in the brain was 40-fold and 7-fold higher at day 4 and 8, respectively (Figure 1). These data clearly show that the brain is copper replete relative to the lung, consistent with previous reporter-based assays. These data are also consistent with the relative expression of *HSP90* from the *pCTR4-2-HSP90* strain (18) in the lung and brain and support the concept that pC*TR4-2* may be applicable to modulating the expression of *C. neoformans* genes during mouse infection.

**Figure 1.**
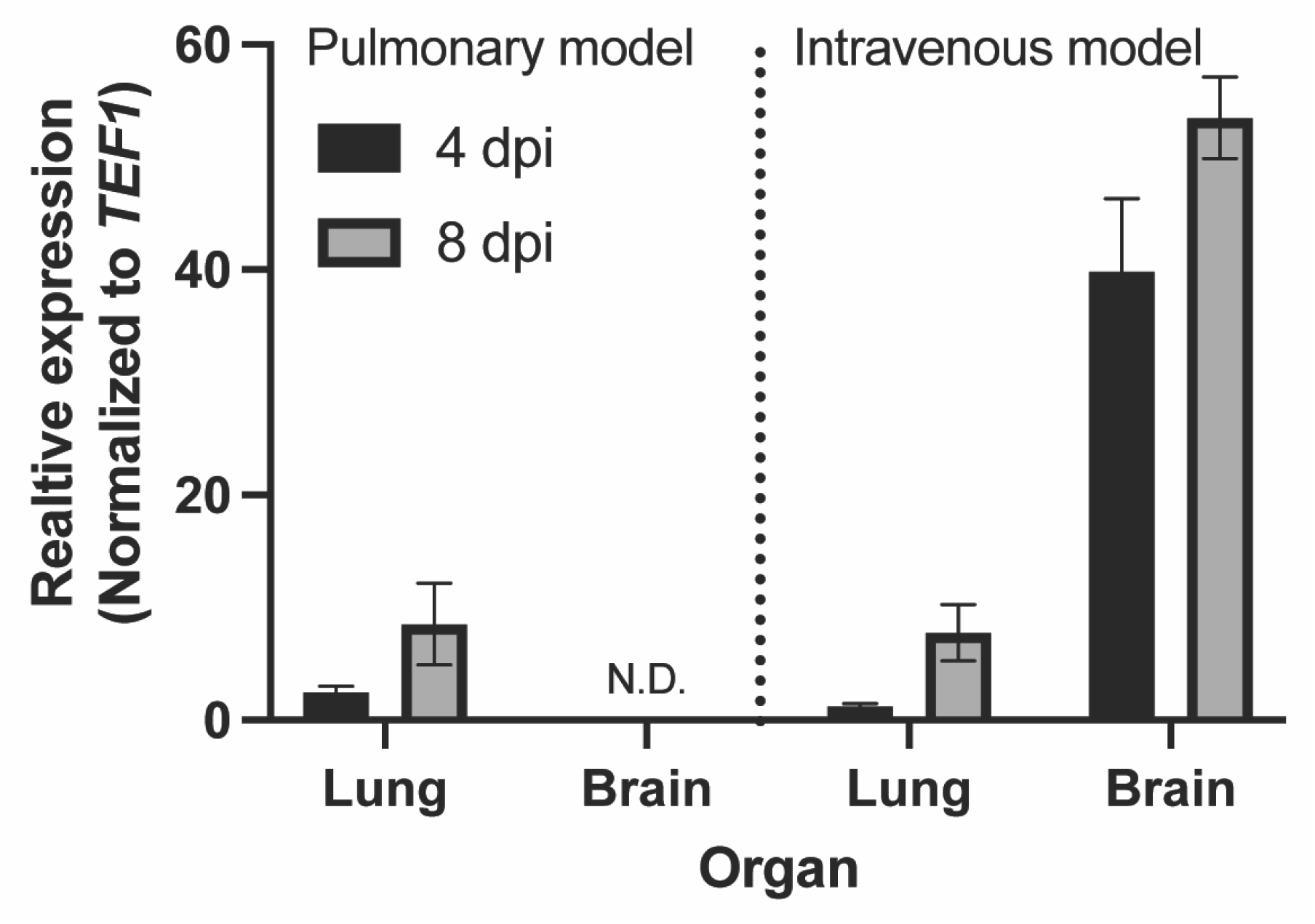
The expression of the copper regulated transporter, *CTR4*, is higher in the brain than in the lungs. Brains and lungs were harvested from mice 4- or 8-days post inoculation (dpi) with KN99α via intranasal instillation (pulmonary model) or lateral tail vein injection (intravenous model). Data represents mean and SEM of three mice per group. N.D.=none detected.

### Application of p*CTR4-2* to the 1,3-β-glucan synthase catalytic subunit, *FKS1*

The echinocandin class of antifungal drugs is the first to directly target the fungal cell wall and is the therapy of choice for invasive *Candida* infections (25). Echinocandins are also active against *Aspergillus fumigatus* and cause lysis of hyphal compartments, leading to fungistatic activity (26). The minimum inhibitory concentration (MIC) of echinocandins against *C. neoformans* is much higher than the MIC for *Candida* spp. or the minimum effective concentration (MEC) for *Aspergillus* spp. A typical MICs for caspofungin against *C. neoformans* is 16 μg/mL while MICs for susceptible *Candida* and *Aspergillus* are 10-1000-fold lower (27). The mechanism for the reduced susceptibility of *C. neoformans* to echinocandins is the subject of active investigation (28–31). Although recent work has shed light onto genes and processes that contribute to the resistance, the fundamental molecular basis of this resistance remains to be described. The *C. neoformans* Fks1 is inhibited by echinocandins *in vitro* (22) and the lack of successful attempts to generate deletion mutants indicate *FKS1* is essential (21). To further confirm the latter finding and to test the effect of reduced *FKS1* expression on the biology of *C. neoformans* and fitness during mouse infection, we constructed a strain containing the p*CTR4-2-FKS1* allele.

To do so, we utilized the recently optimized CRISPR/Cas9 system reported by Huang *et al*. to knock-in the p*CTR4-2* construct immediately upstream of the start codon (32). On YPD and medium containing BCS, two independent strains grew similarly to the H99 parental strain at 37°C (Figure 2A; data not shown). In high copper medium (50 μM CuSO_4_), the p*CTR4-2-FKS1* strain showed a modest reduction in growth relative to the parental strain. The growth defect is less apparent when the strains are incubated at 30°C (Figure 2A) and is not exacerbated with higher copper concentrations (up to 200 μM, data not shown). Elevated temperature causes cell wall stress in *C. neoformans* and activates the cell wall integrity MAP kinase pathway (33). Notably, we observed the appearance of isolated suppressor colonies when the p*CTR4-2-FKS1* strain was grown at 37°C on YPD supplemented with copper, the p*CTR4-2-FKS1* strain (Figure 2A). These larger colonies did not emerge when H99 was incubated under the same conditions. Finally, we tested whether the addition of sorbitol, an osmotic stabilizer, could rescue the growth defect of the p*CTR4-2-FKS1* strain in the presence of copper. Although we did not observe any rescue of the overall growth of p*CTR4-2-FKS1* strain on YPD supplemented with 1M sorbitol and 50 μM CuSO_4_, the generation of suppressors was eliminated (Figure 2B), suggesting that under these conditions, this strain is experiencing significant cell wall stress which is reduced when grown on an osmotic stabilizer.

**Figure 2.**
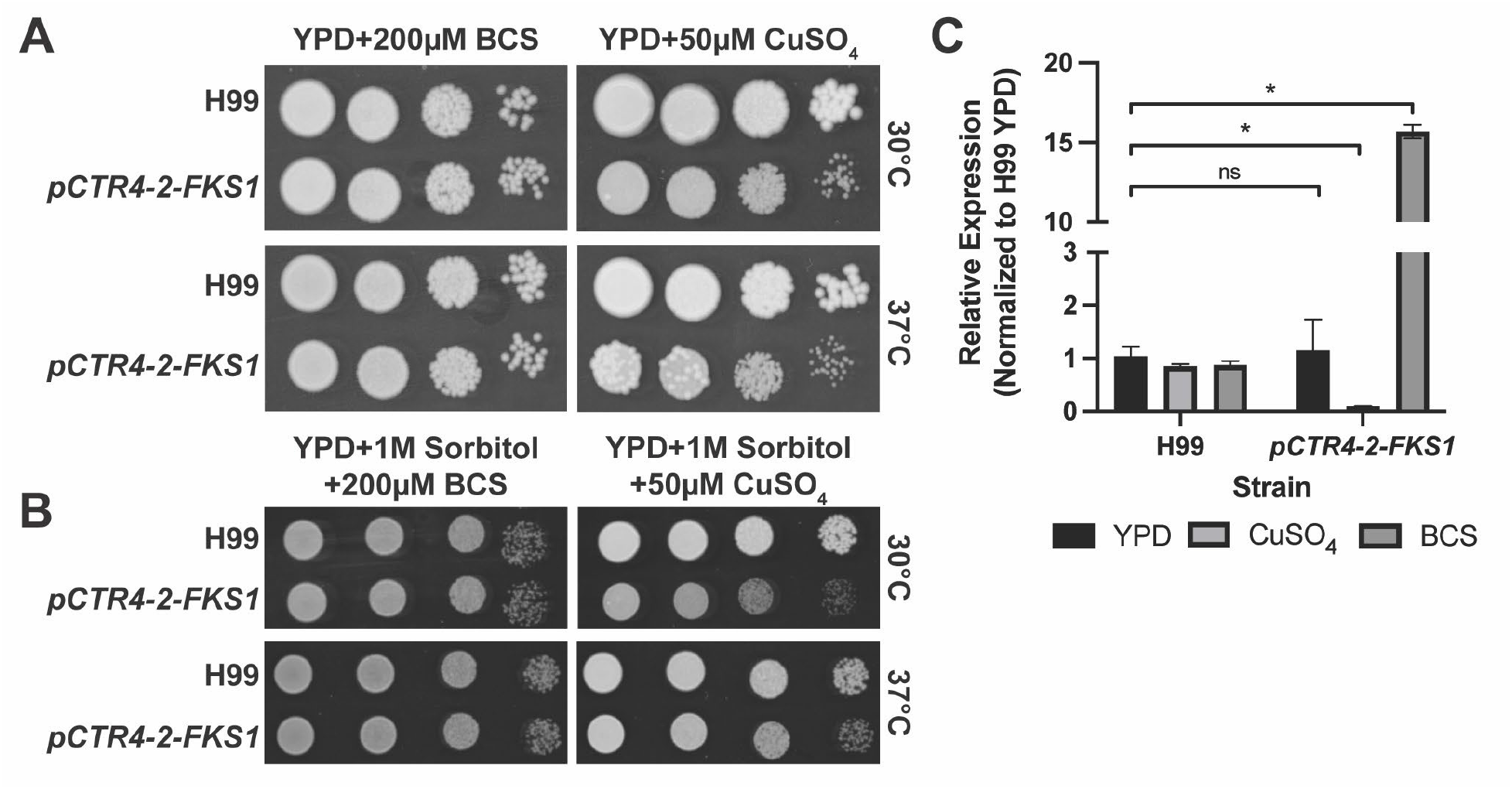
Expression of *FKS1* in *pCTR4-2-FKS1* responds to copper. A) Ten-fold serial dilutions of H99 and *pCTR4-2-FKS1* on YPD (A) or YPD+1M Sorbitol (B) supplemented with 200 μM BCS or 50 μM CuSO_4_. Plates were incubated at 30°C or 37°C for 72 hours. C) Expression of *FKS1* in *pCTR4-2-FKS1* is reduced 10-fold in YPD and with 50μM copper sulfate and is overexpressed with the addition of 200μM BCS. Data represents mean and SEM of 3 biological replicates. *p<0.007; n.s.=not significant by unpaired t-test compared to H99 YPD.

The relative insensitivity of the p*CTR4-2-FKS1* strain to copper addition could be due to intrinsically low expression of *FKS1*, incomplete suppression, or very little *FKS1* expression is required for replication. To determine if any of these explanations were operative, we compared *FKS1* expression in YPD, YPD+BCS and YPD+50 μM CuSO_4_, the same conditions for which the growth assays were performed. As shown in Figure 2C, the expression of *FKS1* is reduced by 100-fold in copper-supplemented medium relative to the parental H99 strain. Conversely, the addition of BCS restored *FKS1* expression to levels 15-fold higher than H99, suggesting that in YPD, the overexpression of *FKS1* does not confer any growth defects at 30°C or 37°C. When compared to our growth assays, these results are striking. Whereas the suppression of other essential genes, such as *FAS1* or *FAS2*, results in virtually no growth on 25 μM copper (15), the suppression of *FKS1* by 100-fold causes only a modest growth defect (Fig. 2A/B). These data suggest that *C. neoformans* can withstand a significant reduction in *FKS1* expression without severe effects on fitness.

### Reduced expression of *FKS1* results in activation of the cell wall integrity pathway and remodeling of the cell wall

Although copper containing media reduces the growth of the p*CTR4-2-FKS1* strain modestly, the overall viability is striking when considering expression of this essential gene is reduced by 100-fold. To determine whether this significant reduction in *FKS1* expression results in a functional reduction of activity, we examined the effect of copper repression on the susceptibility of p*CTR4-2-FKS1* to caspofungin. We performed checkerboard assays with increasing concentrations of caspofungin and copper sulfate (Figure 3A/B). In the absence of copper, the MIC of p*CTR4-2-FKS1* is identical to H99 (Figure 3C). The MIC of H99 is 16 μg/mL and this is unaffected by increasing concentrations of copper (Figure 3A). However, the MIC of p*CTR4-2-FKS1* decreases four-fold to 4 μg/mL in the presence of 40 μM copper sulfate (Figure 3B). Overexpression of *FKS1* with the addition of BCS did not change the caspofungin MIC compared to H99 (Figure 3C), suggesting increasing target abundance is not enough to drive increased caspofungin resistance.

**Figure 3.**
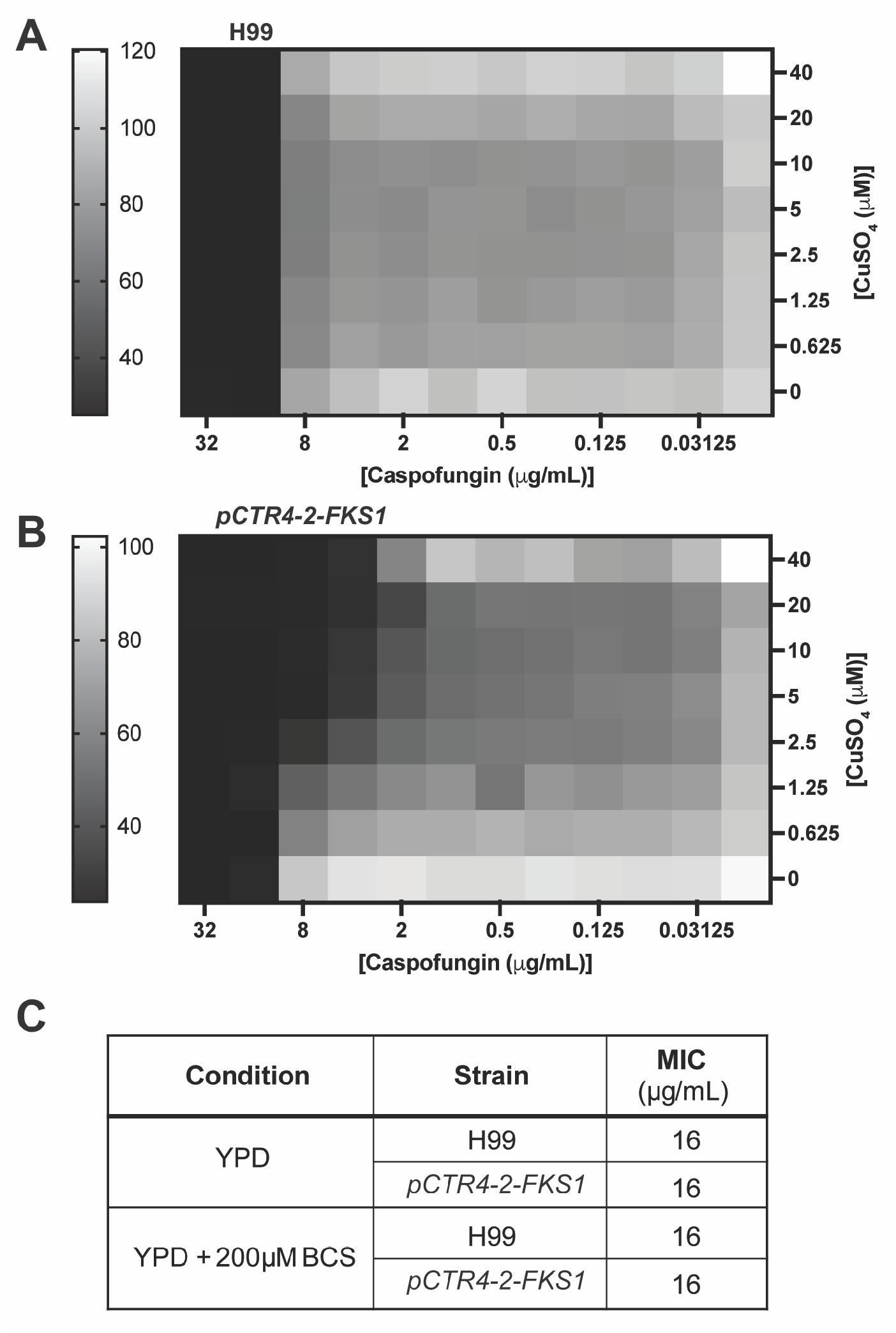
Copper increases the sensitivity of *pCTR4-2-FKS1* to caspofungin. FIC checkerboard assay with H99 (A) or *pCTR4-2-FKS1* (B) with increasing concentrations of caspofungin and copper sulfate. Heatmaps represent percent growth compared to the untreated control as determined by OD_600_ reading after 48 hours at 37°C. Representative plates of two independent replicates. C) Caspofungin MICs in YPD or YPD supplemented with 200μM BCS.

Reduction in Fks1 activity with caspofungin treatment activates of the cell wall integrity pathway (CWIP) resulting in a compensatory increase in chitin (30, 34). Thus, we hypothesized that our p*CTR4-2-FKS1* strain would have a similar response when grown under repressive conditions. To test this, we incubated H99 or p*CTR4-2-FKS1* in YPD, YPD supplemented with 50 μM copper sulfate, 200 μM BCS or a subinhibitory concentration of caspofungin (8 μg/mL), to mid log phase then performed western blots for phosphorylated Mpk1 (pMpk1). Mpk1 is the terminal MAP kinase in the CWIP and is phosphorylated in response to cell wall stresses such as exposure to caspofungin (35). When incubated in either YPD or YPD supplemented with copper sulfate, Mpk1 phosphorylation was increased in p*CTR4-2-FKS1* compared to H99. This increase was even more apparent upon treatment of p*CTR4-2-FKS1* with caspofungin (Figure 4A). In the presence of BCS, the amount of pMpk1 showed by the p*CTR4-2-FKS1* was similar to H99. The correlation between the level of *FKS1* expression and activation of CWIP is consistent with *C. neoformans* mounting a compensatory response to reduced Fks1 activity.

**Figure 4.**
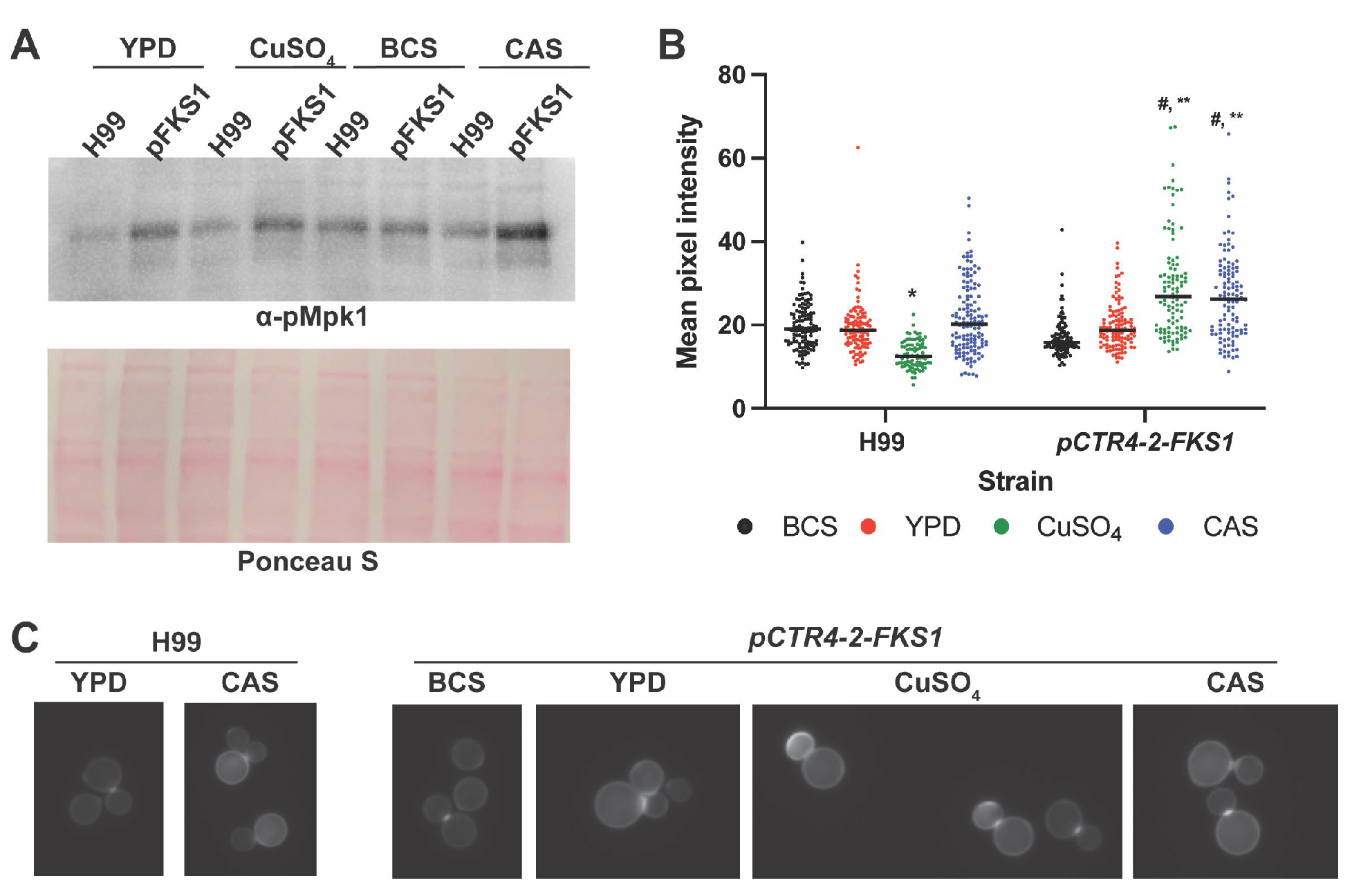
Knockdown of *FKS1* results in activation of the CWIP and an increase in chitin in the cell wall. **A)**Western blotting of pMpk1 (indicated with arrow) after indicated treatments. Ponceau S shown as loading control. B) Mean pixel intensity of CFW stained cells quantified using ImageJ. Representative data from two independent experiments with at least 100 cells per condition. Line represents median value. *,**,^#^ p<0.0001 compared to H99 YPD (*), *pCTR4-2-FKS1* YPD (**) or BCS (#) by one way ANOVA with Tukey’s multiple comparisons. C) Representative images of CFW stained cells. BCS=200μM BCS, CuSO_4_=50μM copper sulfate, CAS = 8μg/mL caspofungin.

One of the components of the cell wall stress response mediated by the CWIP is increased deposition of chitin within the cell wall (36). To determine whether activation of the CWIP in p*CTR4-2-FSK1* results in changes to the cell wall, we stained cells with calcofluor white (CFW) to characterize the amount of cell wall chitin. In general, we observed the typical chitin distribution on H99 cells grown in YPD, YPD+200 μM BCS, and YPD+50 μM copper sulfate where staining is brightest at the septum with less intense but uniform staining of the lateral cell wall (Figure 4B, C). Interestingly, we observed a significant decrease in overall staining intensity upon treatment with 50 μM CuSO_4_. As expected, treatment of H99 with caspofungin resulted in a uniform increase in lateral cell wall staining (Figure 4B/C).

Consistent with our western blot data, we observed similar levels and distribution of chitin between H99 and p*CTR4-2-FKS1* in the presence of BCS. Compared to BCS treatment, chitin levels increased slightly in YPD with a uniform increase in lateral cell wall staining. However, with the addition of 50 μM copper sulfate, chitin staining intensity of p*CTR4-2-FKS1* increased significantly. Furthermore, the distribution of chitin was altered with many cells displaying extremely bright staining of daughter cells or bright, non-uniform patches along the lateral cell wall. Treatment of p*CTR4-2-FKS1* with caspofungin increased chitin compared to YPD alone, however the distribution in these cells more closely resembles YPD with a uniform increase in chitin staining of the lateral cell wall. Together these data are consistent with the conclusion that the copper-induced reduction in the expression of *FKS1* in the *pCTR4-2*-*FKS1* strain results in a functional reduction in Fks1 activity which subsequently activates the CWIP and a compensatory increase in chitin synthesis.

### Altered *FKS1* expression *in vivo* results in reduced fitness in murine model of cryptococcosis

Next, we sought to determine whether the *pCTR4-2* expression system could be used to evaluate the effect of reduced *FKS1* expression on fitness during infection. To test this, we used the intravenous model of cryptococcosis, where both the lungs and the brain become infected rapidly, resulting in reproducible fungal burden in both organs within 4 days. In this way, we could directly compare expression and fungal burden in the organs of the same animal. We inoculated AJ/cr mice with H99 or p*CTR4-2-FKS1* via the lateral tail vein then harvested brains and lungs at 4 days post inoculation. To confirm whether the expression of *FKS1* in p*CTR4-2-FKS1* responds to the copper environment of each organ, we measured *FKS1* expression in both the lungs and the brains. In accordance with *CTR4* expression data (Figure 1), the expression of *FKS1* in the lungs was significantly reduced (p=0.0022 by Mann-Whitney test) in p*CTR4-2-FKS1* compared to H99 (Figure 5A). The reduction in *FKS1* expression *in vivo* was not as pronounced as observed *in vitro* (Figure 2C); but was consistent with the expression of *CTR4 in vivo* (Figure 1). Conversely, *FKS1* expression in the brain was significantly increased in the in p*CTR4-2-FKS1* strain relative to H99 (Figure 5A; p=0.0022 by Mann-Whitney test) which, again, follows the trend observed for *CTR4* expression (Figure 5A). These data are consistent with the notion that the lungs are copper-replete while the brain is copper limited and the copper-regulatable promoter can be used to modulate the expression of genes in murine models of infection.

**Figure 5.**
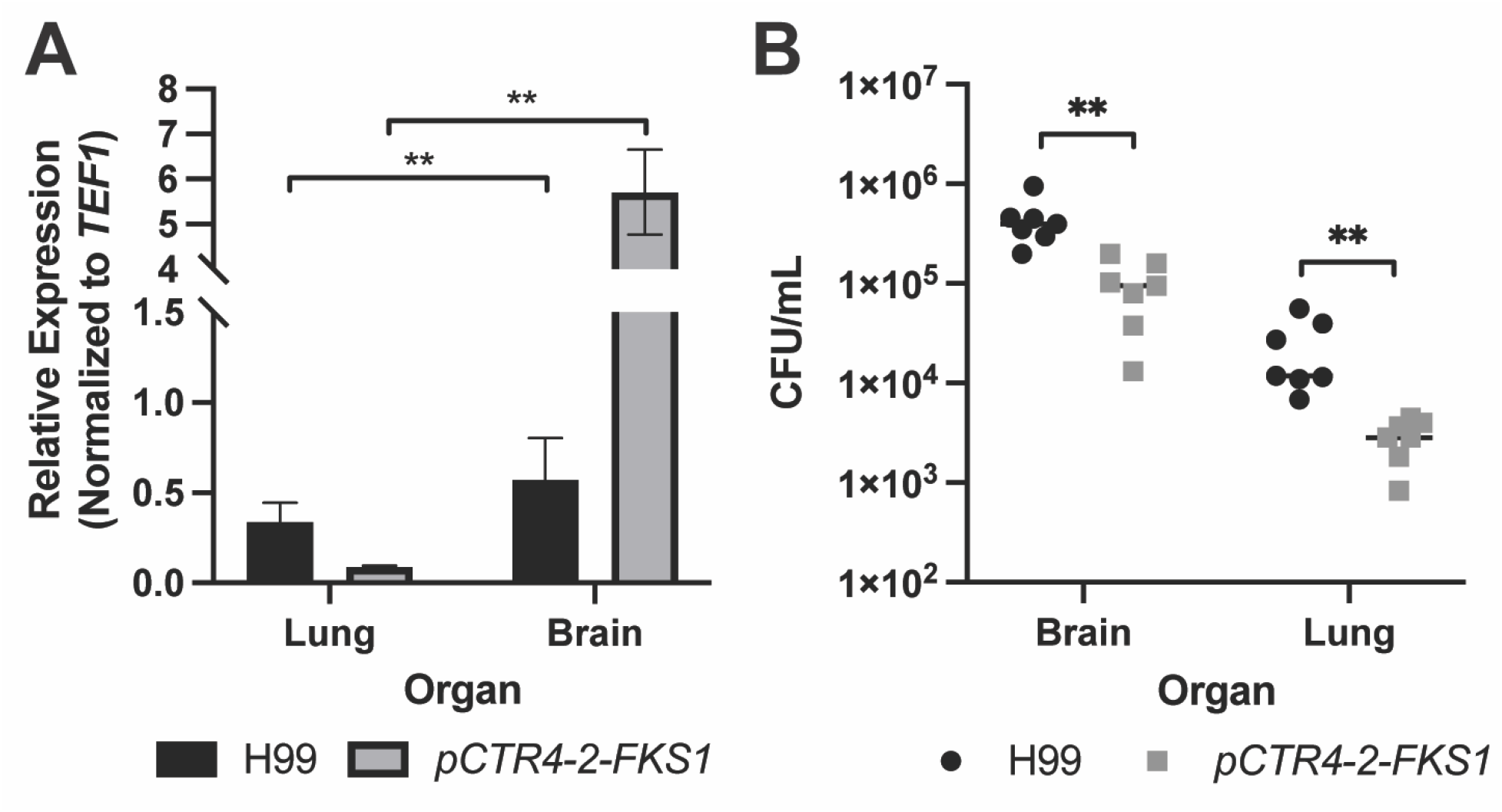
The repression or overexpression of *FKS1 in vivo* results in a fitness defect. A) Expression of *FKS1* in the lungs and brains collected 4 dpi from mice inoculated via lateral tail vein with H99 or *pCTR4-2-FKS1*. Data represents mean and SEM of *FKS1* expression normalized to *TEF1*. n=6 mice per group. **p=0.0022 by Mann-Whitney test comparing *pCTR4- 2-FKS1* to H99 for each organ. B) Fungal burden of lungs and brains of mice collected 4dpi. Data represent individual values of 7 mice per group. **p<0.001 by Mann-Whitney test comparing H99 to *pCTR4-2-FKS1* in each organ corrected for multiple comparisons with a Bonferroni correction.

Finally, we measured the fungal burden in lung and brain tissue of mice inoculated with H99 or p*CTR4-2-FKS1* to determine if modulation of *FKS1* expression *in vivo* results in a fitness cost. In the lungs, where *FKS1* expression is significantly reduced in p*CTR4-2-FKS1* compared to H99, we observed a ~1 log_10_ reduction in fungal burden (Figure 5B). Despite the expression of *FKS1* in the p*CTR4-2-FKS1* being much closer to H99 in vivo compared to in vitro, the fitness defect is much more pronounced in vivo. This suggests that maintenance of *FKS1* activity is much more important for *C. neoformans* survival in vivo than in vitro.

While overexpression of *FKS1 in vitro* had no effect on fitness, the fungal burden of brains inoculated with p*CTR4-2-FKS1* was reduced by about one-half log_10_ compared to H99 (p=0.0012 by Mann-Whitney test), suggesting that overexpression of *FKS1 in vivo* is detrimental to *C. neoformans* growth or survival in the brain (Figure 5B). Interestingly, we observed the same results with control experiments performed with the p*CTR4-2-FAS1* strain studied by the Perfect lab; specifically, overexpression of *FAS1* reduced fungal burden in the brain (Figure S1).

The Perfect lab also reported that the brain burden for this strain was reduced after dissemination from the lung. Together, these data support the hypothesis that the regulation of *FKS1* expression is required for full fitness *in vivo* and that reduced *FKS1* activity is more important for fitness during infection than under standard *in vivo* conditions.

## Discussion

The goals of this study were two-fold: 1) explore the scope of copper-regulated gene expression as an approach to studying essential genes in *C. neoformans* infection models and 2) characterize the effect of reduced expression of the putatively essential gene *FKS1 in vitro* and *in vivo*.

To our knowledge, this is the fourth gene that has been studied *in vivo* using the *CTR4-2* promoter to modulate expression (15, 18). Although this approach has utility, there are some important limitations to the system that must be considered during experimental design and data interpretation. First, this system should always be evaluated on a gene-to-gene basis. For any given gene, the wild type levels of expression and the amount of repression needed to see a phenotype can vary. For example, a gene with very low expression may already be expressed at levels similar to *CTR4* expression in copper-replete conditions and may not be repressed *in vivo* to the point where robust phenotypes can be detected. Conversely, highly expressed genes may still be “repressed” under copper depleted conditions; in such cases, the p*CTR4-2*-regulated allele will represent a knock-down in both conditions. Second, it is critical that the expression of the target gene be checked *in vitro* and *in vivo* to ensure that gene expression is responding to copper concentration. When manipulating essential genes, chromosomal rearrangements and/or mutations can occur during strain generation or as the strain is passaged in routine lab use. These alterations could result in unexpected expression patterns. Third, because the copper concentrations cannot be changed in the animal, this promoter system is limited to the study of the effect of reduced gene expression during the establishment and progression of pulmonary infection, or in some cases, to assess the effects of gene overexpression in the brain. With these limitations in mind, however, this system can be a powerful tool for the evaluation of essential *C. neoformans* genes *in vivo*.

We chose to further explore the *in vivo* utility of the *pCTR4-2* promoter system by applying it to β-1,3-glucan synthase, *FKS1*, to learn more about the effect of reduced expression on *C. neoformans in vitro* and *in vivo*. As briefly outlined in the introduction, *C. neoformans* is notably resistant to echinocandins with MIC values orders of magnitude higher than other pathogenic yeasts. Previous genetic experiments indicate that *FKS1* is essential and that the enzyme, Fks1, is inhibited by echinocandins in a manner similar to enzymes isolated from susceptible species (22, 37). Gold particle immune-electron microscopy studies with antibodies specific to 1,3-β-glucan linkages indicate that *C. neoformans* cells contain this polymer and that treatment with an echinocandin reduces their levels in the cells (36). Consistent with other yeasts, exposure of *C. neoformans* to echinocandins also activates the cell wall stress response mediated by the CWIP (Figure 4A; (35)). The strong correlation between the effects of echinocandins on susceptible yeast species and *C. neoformans* suggests that the mechanism of action of the drug is similar amongst the discordant organisms.

Our studies with the p*CTR4-2-FKS1* strain make further contributions to our understanding of the phenotypes associated with reduced glucan synthase expression. First, suppression of *FKS1* expression leads to activation of the CWIP as determined by increased phosphorylation Mpk1 under those conditions. This activation is seen in other pathogenic fungi as well (38). Second, suppression of *FKS1* expression increases susceptibility of the strain to caspofungin, although it was reduced by only 4-fold despite a 100-fold reduction in *FKS1* expression. In *C. albicans*, overexpression of *FKS1* did not increase the MIC of caspofungin toward planktonic cells; it did, however, increase the MIC in *C. albicans* biofilms (39). Third, cells treated with either caspofungin or that have copper-reduced expression of *FKS1* show increased chitin within the cell wall (Figure 4C/D). Chitin has been shown to be a key modulator of echinocandin tolerance and resistance (40). Fourth, the additional of an osmotic stabilizer (1M sorbitol) to the medium reduces the effect of reduced *FKS1* expression on growth. This phenomenon is well-described for a range of cell wall-targeting agents and fungal pathogens (36). Thus, the phenotypes associated with the genetic reduction in *FKS1* expression correlate quite well with the effects of caspofungin exposure in other yeast.

A key finding of our work is that the *C. neoformans* tolerates a dramatic reduction in *FKS1* expression without significant changes in growth/viability *in vitro*. This raises the question what mediates this tolerance? First, it is possible that only a very small amount of transcript is required to maintain 1,3-β-glucan synthase protein at levels needed for preserve viability. Kalem *et al*. has shown that the regulation of *FKS1* mRNA stability and translation plays an important role in resistance to caspofungin (31). Second, it is possible that the Fks1 protein is remarkably stable and there is little correlation between gene and protein expression. If the difference in expression were only a few-fold then this would seem much more likely explanation; since the difference is two orders of magnitude, it is very difficult to invoke this mechanism. Again, Kalem *et al*. found that that there was only a ~1.5-fold difference between mRNA and Fks1 protein levels in a mutant with increased translation of *FKS1* message (31).

A third explanation for our observations is that the cell wall stress-induced compensatory response triggered by reduced Fks1 activity in *C. neoformans* is much more effective than for other organisms. A substantial number of genes not directly related to glucan synthesis have been shown to play important roles in *C. neoformans* resistance to echinocandins (28–31). These may combine with the CWIP-mediated responses to effectively buffer the cell against reduced Fks1 activity/*FKS1* expression in a manner not observed for other yeasts. This would explain the apparent disconnect between the essential nature of the *FKS1* gene (21) and the tolerance of *C. neoformans* to all but a, presumably, near complete depletion/inhibition of 1,3-β-glucan synthase activity. Kraus et al. have suggested that the calcineurin and CWIP effectively compensate for reduced 1,3-β-glucan synthase activity in the absence of other stressors (35). As part of their studies, Kraus et al. found that *FKS1* expression is induced by activation of the CWIP. Our data suggest that increased expression of *FKS1* is unlikely to have a significant role in the response to reduced 1,3-β-glucan synthase activity and that other mechanisms such as altered chitin homeostasis appear to be more important. The altered chitin content of strains with reduced *FKS1* expression and the role of chitin in the response to lowered 1,3-β-glucan synthase activity further support that hypothesis.

Interestingly, we observed that treatment of H99 with 50 μM copper sulfate results in a significant reduction in overall chitin levels. Links between the cell wall and copper homeostasis have been postulated previously, as the copper responsive transcription factor Cuf1 regulates cell wall synthesis genes (41). Recently, Probst *et al*., solidified a connection between copper homeostasis and the cell wall, showing that mutation of copper homeostasis machinery results in changes to the levels of chitin in the cell wall (42). Similar to our results reported here, Probst *et al*. show calcofluor white images in copper sufficiency with less intense staining than those in copper-deficient media.

We also tested the ability of this strain to establish infection in a murine model of cryptococcosis. In both the brain and lungs, the modulation of expression of *FKS1 in vivo* by copper resulted in significant reduction in fungal burden. As we expected, in the copper-replete lungs, the expression of p*CTR4-2-FKS1* was significantly reduced resulting in a ~1 log_10_ reduction in fungal burden compared to animals inoculated with H99. In the copper-deplete brain, the expression of *FKS1* was significantly increased in p*CTR4-2-FKS1* compared to H99, however we observed a half-log reduction in fungal burden. These results support the idea that proper regulation of *FKS1* and the cell wall composition is critical for full fitness *in vivo*. Although we did not see any associated defects *in vitro*, the environment *in vivo* is much more complex and dynamic. For example, copper is subject to regulation by the host as part of nutritional immunity (43), thus copper concentrations within the microenvironment of fungal lesions can change rapidly. In *M. tuberculosis*, copper has been shown to accumulate at granulomatous lesions in the lung (44). In addition, macrophages use copper as an antimicrobial by pumping high levels of copper into phagolysosomes, resulting in higher copper exposure to cells within macrophages. In addition, even slight changes to the fungal cell wall can have profound impacts on disease pathology given that the cell wall is the interface between fungal and host cells, and perturbations in cell wall architecture often leave pathogen recognition patterns exposed.

Finally, we note that reduction in *FKS1* expression by ~4-fold results in a 1 log_10_ reduction in fungal burden while in vitro the growth is only minimally affected. This suggests that the compensatory mechanisms that support the ability of *C. neoformans* to tolerate reduced 1,3- β-glucan synthase expression are much less effective *in vivo*. One explanation of this finding is that *C. neoformans* cells are subject to a combination of stressors that all must be managed through the same compensatory mechanisms. Although further work will be required to understand the contributing factors and mechanisms, it is clear that 1,3-β-glucan synthase expression is much more important *in vivo* than would be predicted from in vitro phenotypes.

This work demonstrates the utility of the copper-regulated promoter, p*CTR4*, in assessing the role of essential genes in animal models of infection, a critical step in target validation for the development of novel antifungal drugs. Furthermore, we have begun to further unravel the mystery of caspofungin resistance in *C. neoformans*. Our data support the idea that *C. neoformans* can tolerate a significant reduction in Fks1 activity, resulting in reduced efficacy of caspofungin despite direct inhibition of the enzyme itself. A complete understanding of how the cell wall architecture and response to cell wall stress differs between *C. neoformans* and other pathogenic fungi can inform the development of cell wall active antifungals and understanding how *C. neoformans* interfaces with the host during disease.

## Materials and Methods

### Strains, media and chemicals

All experiments were performed with *C. neoformans* H99 Stud. *C. neoformans* strains were maintained on YPD and stored at −80°C in 25% glycerol. Plasmid pCTR4-2 was generously provided by Dr. Tamara Doering (Washington University, St. Louis MO). Plasmids BHM2329 and BHM2403 were generously provided by Dr. Hiten Madhani (University of San Francisco, San Francisco CA). Copper sulfate pentahydrate (Cat# C7631), bathocuproinedisulfonic (BCS; Cat#146625) acid disodium salt hydrate, and caspofungin diacetate (Cat# SML0425) were obtained from Sigma.

### Construction of plasmids

To make pUL-HYG, pZY97 (45) was digested with SacI and AatII (New England Biolabs, Cat# R0157 and R0117) simultaneously to drop out the existing resistance cassette and generate the cut vector for Gibson assembly. The hygromycin resistance (HYG^R^) cassette was amplified from pXL1-pTEF1 (46) with primers LCR033 and LCR012 designed to overlap the cut vector by 20bp to facilitate Gibson assembly and add a 20bp universal linker at each end of the cassette. 100ng of cut vector with a 2-fold molar excess of insert were combined in the Gibson assembly reaction (New England Biolabs, Cat# E5510S), following manufacturer instructions. Plasmids were isolated from individual clones and the universal linker region was confirmed at each end of the resistance cassette by Sanger sequencing with primers LCR033 and LCR034. Functionality of the inserted resistance cassette was confirmed by transformation into H99 and selection on 400mg/ml hygromycin plates.

We generated plasmid pSB-CTR4 containing the pCTR4-2 promoter downstream of HYG^R^ to facilitate promoter replacement of additional genes by simply amplifying the resistance cassette and pCTR4-2 promoter with primers containing short-arm homology to a gene of interest. The promoter replacement construct was generated by amplifying the pCTR4-2 promoter from plasmid pCTR4-2 (17) with primers SP181 and SP196 to add homology to the HYG^R^ cassette at the 5’ end of the product and homology to pUC19 at the 3’ end of the product. The HYG^R^ cassette was amplified with primers SP197 and SP180 from pUL-HYG. The 5’ end of this product contains homology to pUC19. These fragments were cloned into pUC19 with the HYG resistance cassette upstream of pCTR4-2 with InFusion cloning (Takara, Cat#638911). Plasmids were isolated from individual clones and confirmed by digestion with HindIII (New England Biolabs, Cat# R0104) and the resulting plasmid pSB-pCTR4 was used to amplify the final *FKS1* promoter replacement cassette.

### Construction of pCTR4-2-FKS1

pCTR4-2-FKS1 was generated using CRISPR-Cas9 with short-homology repair (32). CnoCas9 was amplified using SP29 and SP30 from BHM2403 (32). The sgRNA was designed using the Eukaryotic Pathogen CRISPR guide Design tool (47). To generate the complete sgRNA construct, SP106 and SP200 and SP105 and SP201 were to amplify the U6 promoter and sgRNA scaffold, respectively with the 20nt guide sequence, from plasmid BHM2329. These two fragments were joined with overlap-extension PCR with primers SP31 and SP32.

The promoter replacement construct was amplified from pSB-pCTR4 with primers SP199 and SP202 containing 50bp of microhomology to the *FKS1* 5’ UTR. Electroporation-mediated transformation was used to transform 1μg Cas9, 1μg gRNA and 3μg of *pCTR4-2-FKS1* promoter construct (48) and cells were plated on YPD with 400μg/mL hygromycin B (Alfa Aesar, Cat# J69681). Transformants were verified with PCR spanning the joint between the promoter construct and *FKS1* coding sequence and tested for response to copper by growth assay and gene expression.

### Antifungal susceptibility and interaction assays

Minimum inhibitory concentrations were determined using CLSI guidelines (49). Yeasts were cultured overnight in 3ml YPD at 30°C, then washed twice in sterile PBS. Two-fold serial dilutions of each drug were prepared in RMPI+MOPS pH 7 (Gibco RPMI 1640 with L-glutamine [Cat# 11875-093] and 0.165M MOPS), then 1×10^3^ cells were added per well in a total volume of 200μL. Plates were incubated at 37°C for 72 hours.

For fractional inhibitory concentrations assays, standard checkerboard assays were performed as previously described (50). Briefly, two-fold dilutions of caspofungin or CuSO_4_ were prepared in YPD at four-fold the desired final concentration. 50μL of the caspofungin dilution series was dispensed into a 96-well plate with the concentration decreasing across the columns. Next, 50μL of the CuSO_4_ dilution series was dispensed into the plate with the concentration decreasing down the rows, such that the top corner of the plate contained the highest concentration of each compound and the opposite corner contained vehicle only. 1×10^3^ cells were added per well for a total volume of 200μL. Plates were incubated for 42 hours at 37°C, then MIC of each drug alone, and each drug in combination was determined.

### Plate-based growth assay

Overnight cultures of H99 and *pCTR4-2-FKS1* were diluted to OD_600_=1 then ten-fold serial dilutions were prepared in PBS. 5μL of each dilution was spotted on indicated media, then incubated for 48-72 hours at 30°C or 37°C. Plates were imaged using an Epson Perfection V600 Photo Scanner. Contrast was adjusted equally across all images for the best visualization of colonies.

### Characterization of Mpk1 phosphorylation by western blot

Overnight cultures of H99 and pCTR4-FKS1 were diluted to OD=0.1 in YPD, YPD + 50μM CuSO4, or YPD + 200μM BCS then grown to mid-log phase (4 hours) at 30°C with shaking at 200rpm. Protein was extracted in extraction buffer (10 mM HEPES pH 7.4 to 7.9, 1.5 mM MgCl_2_, 10 mM KCl, 1 mM dithiothreitol [DTT], 1× HALT protease and phosphatase inhibitor cocktail (Thermo Scientific Cat #1861280)) by five bead beating cycles of 30 seconds followed by 30 seconds on ice per cycle. Debris and beads were pelleted before supernatant was recovered and the protein concentration was quantified by Bradford assay. Protein (20μg) was loaded on a 10% SDS-PAGE gel and run at 80V. Samples were transferred to nitrocellulose membrane for 1 h at 100V, then membrane was stained with Ponceau for 5 min at RT. The membrane was blocked with 5% bovine serum albumin (BSA) in Tris-buffered saline with Tween-20 (TBST) for 1 hour at RT, then the membrane was incubated with 1:2,000 rabbit anti-p-p44 (Cell Signaling, Phospho-p44/42 MAPK, Cat# 4370) in 5% BSA/TBST overnight at 4°C. The membrane was washed 3 times for 5 minutes with TBST, then incubated for 1 hour at RT with 1:10,000 goat anti-rabbit horseradish peroxidase (HRP) (Bio-Rad, Cat# STAR208P). The membrane was washed 3 times for 5 minutes with TBST, then developed with chemiluminescent substrate and imaged on a Thermo Scientific My ECL imager.

### Microscopy-based characterization of chitin content and distribution

Cells were cultured under the same conditions as above. For calcofluor white (CFW), cells were washed twice with PBS, then resuspended in 10μg/mL CFW in PBS (Fluorescent brightener 28, Sigma, CAT#F3543) and incubated for 20 minutes at RT in the dark. Cells were imaged on a Nikon epifluorescence microscope with a Cool Snap HQ2 camera and Nikon Elements image acquisition and analysis software. Mean pixel intensity of images was measured using ImageJ. Images included in the figure were processed in Photoshop only to increase ease of viewing. All images were adjusted equally.

### *In vitro* quantitative RT PCR

Overnight cultures of H99 and pCTR4-FKS1 were diluted to OD=0.1 in YPD, YPD + 50μM CuSO4, or YPD + 200μM BCS then grown to mid-log phase (4 hours) at 30°C with shaking at 200rpm. Cells were collected, pellets were frozen at −80°C then lyophilized. Dried tissue was homogenized with zircon silica beads using the MP Biologicals Fast-Prep24 homogenizer (MP Biomedicals, Cat#116004500). RNA was isolated using the Pure Link RNA kit (Invitrogen, Cat# 12183025) according to the manufacturers protocol then 5μg of RNA was treated with TURBO DNA-free kit (Invitrogen, CAT#AM1907). 500ng of RNA was used for cDNA synthesis with iScript cDNA synthesis kit (BioRad, Cat# 1708840) and cDNA was diluted 1:5 for qRT-PCR. 2μL of diluted cDNA was used per reaction with iQ SYBR Green Supermix and 0.20μM primers. qRT-PCR was performed on the BioRad CFX Connect using a 3-step amplification with 54°C annealing temperature and melt curve analysis. Primers are listed in Table S1.

### *In vivo* quantitative PCR

Female A/J mice (6 weeks old; Jackson Laboratory) were inoculated with 5×10^4^ CFU/mL in 200μL PBS via the lateral tail vein or 1×10^6^ CFU/mL in 50μL PBS via intranasal instillation. Brains and lungs were collected at 4- and 8-days post inoculation then lyophilized. Tissue was homogenized with 2.3mm zirconia-silicate beads using the MP Biologicals Fast-Prep24 homogenizer (MP Biomedicals, Cat#116004500). 1mL of TRIzole (Invitrogen, Cat#15596026) was added to each sample and incubated at RT for 10 minutes. Samples were clarified by spinning at 10Kxg for 5 minutes, then supernatant was transferred to a new tube and 200μL of chloroform was added with mixing by inversion. Samples were spun at 12Kxg for 15 minutes then aqueous phase was transferred to gDNA removal column (Qiagen RNeasy Plus Mini kit, Cat# 74136) then RNA was purified following manufacturer’s instructions. 500ng of RNA was used for cDNA synthesis with iScript cDNA synthesis kit (BioRad, Cat# 1708840) and cDNA was diluted 1:1 for qRT-PCR. 2μL of diluted cDNA was used per reaction with iQ SYBR Green Supermix and 0.20μM primers. qRT-PCR was performed on the BioRad CFX Connect using a 3-step amplification with 54°C annealing temperature. We observed no amplification with our primer sets on cDNA prepared from mice inoculated with PBS. Primers used are listed in Table S1.

### Determination of fungal burden in mouse model of disseminated cryptococcosis

Female A/J mice (6 weeks old; Jackson Laboratory) were inoculated with 5×10^4^ CFU/mL in 200μL PBS via the lateral tail vein. Brains and lungs were collected at 4-days post inoculation and homogenized in PBS with a benchtop homogenizer (VWR). Homogenates were diluted in a 10-fold dilution series and each dilution was plated on YPD. Plates were incubated at 30°C for 48 hr and colonies were counted. Each group contained 7 mice per group. Differences in fungal burden between groups was analyzed using the Mann-Whitney test in GraphPad Prism 9.

### Ethics Statement

The Guide for the Care and Use of Laboratory Animals of the National Research Council was strictly followed for all animal experiments. The animal experiment protocols were approved by Institutional Animal Care and Use Committee at the University of Iowa (protocol: 7102064).

## Acknowledgments

We thank Xiaorong Lin (Georgia) for helpful discussions. We thank Manning Huang and Hiten Madhani (UCSF) for providing plasmids for *C. neoformans* CRISPR/Cas9-mediated gene eding prior to publication. This work was supported by NIH grants 5R01AI147541 (DJK), T32AI007511 (AJJ), and 5F32AI145160 (SRB).

---

**Figure S1. The fungal burden of p*CTR4-2-FAS1* is significantly lower in both the brain and the lung during infection.** Fungal burden of brains and lungs of mice collected 4 days post-inoculation via the lateral tail vein. Data represent 7 mice per group; control (H99) data is the same data shown in Figure 4A. **p=0.0006 by Mann-Whitney test corrected for multiple comparisons with a Bonferroni correction.

**Table S1. Primers used in this study**

